# SpaTranslator: A deep generative framework for universal spatial multi-omics cross-modality translation

**DOI:** 10.1101/2025.11.15.688644

**Authors:** Hongyu Dong, Sheng Mao, Yukuan Liu, Tian Tian, Lihua Zhang, Juanshu Wu, Shichen Zhang, Peng Jiang, Danqing Yin, Xudong Xing, Peizhuo Wang, Han Li

## Abstract

Recent advances in spatial omics technologies have enabled the simultaneous analysis of multiple molecular patterns in tissue sections, offering unprecedented insights into cellular microenvironments. However, the high cost of measurements and the sparsity of data restrict the availability of paired spatial multi-omics datasets. Here, we present SpaTranslator, a deep generative framework that integrates graph neural networks with an adversarial variational generative model to fully capture spatial characteristics and enable effective cross-modality translation of spatial omics data, enabling simulation of paired spatial multi-omics data from single-omics measurements. Extensive experiments demonstrate that SpaTranslator consistently outperforms baseline methods in both clustering accuracy and biological coherence across various real-world scenarios, including spatial transcriptomics-epigenomics and spatial transcriptomics-proteomics translation tasks. Furthermore, SpaTranslator provides biologically meaningful insights through marker genes and proteins recovery, motif enrichment analysis, and gene regulation inference. Our work offers an effective and adaptable solution for spatial multi-omics cross-modality translation, supporting a broad range of biological and biomedical research.

## 1 Introduction

In recent years, spatial omics technologies have undergone significant development, offering deeper insights into the cellular architecture of relevant tissues. Notable breakthroughs, such as DBiT-seq [1], spatial-CITE-seq [2], spatial ATAC-RNA-seq [3], and MISAR-seq [4], enable simultaneous profiling of multiple spatial omics modalities on a single tissue slice. These techniques concurrently capture multi-modal information for each spot within its spatial microenvironment, thereby enhancing our understanding of the cellular spatial characteristics. However, high measurement costs currently limit the availability of spatial multi-omics datasets. Furthermore, inherent limitations of spatial multi-omics data, such as sparsity and high dropout rates, further restrict the broader applications of these technologies.

With the increasing prevalence of spatial single-omics technologies, such as spatial transcriptomics [5], spatial epigenomics [6], and spatial proteomics [7], single-modality spatial omics data are rapidly accumulating. This presents a valuable opportunity to employ generative frameworks to infer missing modalities for single-modality inputs, thus synthesizing paired spatial multi-omics data. Utilizing generative models to simulate paired spatial multi-omics data *in silico* can substantially reduce experimental costs while increasing data richness and diversity. Notably, large-scale spatial multi-omics data can underpin the development of AI virtual cell foundation models [8], opening new avenues for virtual drug screening, disease process simulation, and precision medicine.

While several existing approaches have been developed for cross-modality translation of single-cell omics data, such as multiDGD [9], scPair [10], JAMIE [11], and scButterfly [12], these methods possess limitations preventing their direct application to spatial omics. First, due to technical constraints, each spot in spatial omics data contains limited molecular information, making it sparser than single-cell data. As a result, integration of information from spatially neighboring spots is often required for downstream analyses [13]. Existing single-cell cross-modality generation approaches generally focus only on feature mapping for individual cells, neglecting spatial neighborhood relationships and thus performing poorly on spatial omics data. Second, most single-cell cross-modality generation algorithms train models on one dataset and directly apply them to others during cross-dataset generation, without considering the inter-dataset heterogeneity (e.g., batch effects). However, there are significant batch effects between spatial omics slices [14]. Existing algorithms that ignore batch effects cannot be effectively applied to spatial cross-modality generation tasks. Finally, many existing approaches are restricted to predefined modality pairs (e.g., transcriptome-to-proteome translation), limiting flexibility as new modalities emerge. With the emergence of new spatial omics sequencing technologies, there is an urgent need for a universal computational method that can effectively address the challenges of data sparsity and batch effects inherent in spatial omics data, and be capable of cross-modality generation across multiple modalities.

In this work, we propose SpaTranslator, a deep generative framework for effective cross-modality translation of spatial multi-omics data. To the best of our knowledge, this is the first approach capable of handling multiple spatial omics modality pairs, including spatial transcriptome–epigenome and spatial transcriptome–proteome translations. Moving beyond existing modality translation methods developed for single-cell omics, SpaTranslator incorporates several components specifically tailored to the unique characteristics of spatial omics. Specifically, SpaTranslator first pre-trains graph neural network (GNN)-based autoencoders to learn informative latent embeddings for each spot by aggregating information from its neighboring spots, thereby effectively mitigating the high sparsity inherent in spatial omics data and fully capturing the spatial characteristics. Additionally, a contrastive learning strategy is employed during pre-training to remove the batch effects between spatial slices. After pre-training, an adversarial variational generative model is introduced to cross-align the latent embeddings derived from the pre-trained autoencoders to enable the cross-modality generation.

Comprehensive evaluation experiments were conducted under diverse scenarios. The results showed that SpaTranslator consistently outperformed baseline methods in the cross-modality translation of spatial multi-omics data under intra-slice and cross-slice translation scenarios, in terms of both quantitative clustering accuracy and qualitative spatial architecture reconstruction. Analyses of the spatial distributions of marker genes and proteins, as well as motif enrichment, validated that SpaTranslator faithfully recapitulated biological features present in real data. Most importantly, the results demonstrated that SpaTranslator enabled the reliable inference of missing spatial assay for transposase-accessible chromatin using sequencing (ATAC-seq) profiles from unimodal spatial transcriptomics, facilitating integrated multi-omics analyses and providing valuable insights into gene regulation and developmental processes from a spatial multi-omics perspective.

## 2 Results

### 2.1 Overview of SpaTranslator

SpaTranslator is a deep generative framework that integrates GNNs with an adversarial variational generative model for cross-modality translation of spatial omics (Fig. 1; Methods). To illustrate, consider the translation from spatial RNA sequencing (RNA-seq) data to spatial ATAC-seq data. Given a reference slice with paired spatial RNA-seq and ATAC-seq data (denoted as *S*1_*A*_ and *S*1_*R*_, respectively) and a target slice with only spatial RNA-seq data (denoted as *S*2_*R*_), SpaTranslator leverages the paired reference data to infer the missing ATAC-seq modality for the target slice (denoted as *S*2_*A*_).

**Fig. 1.**
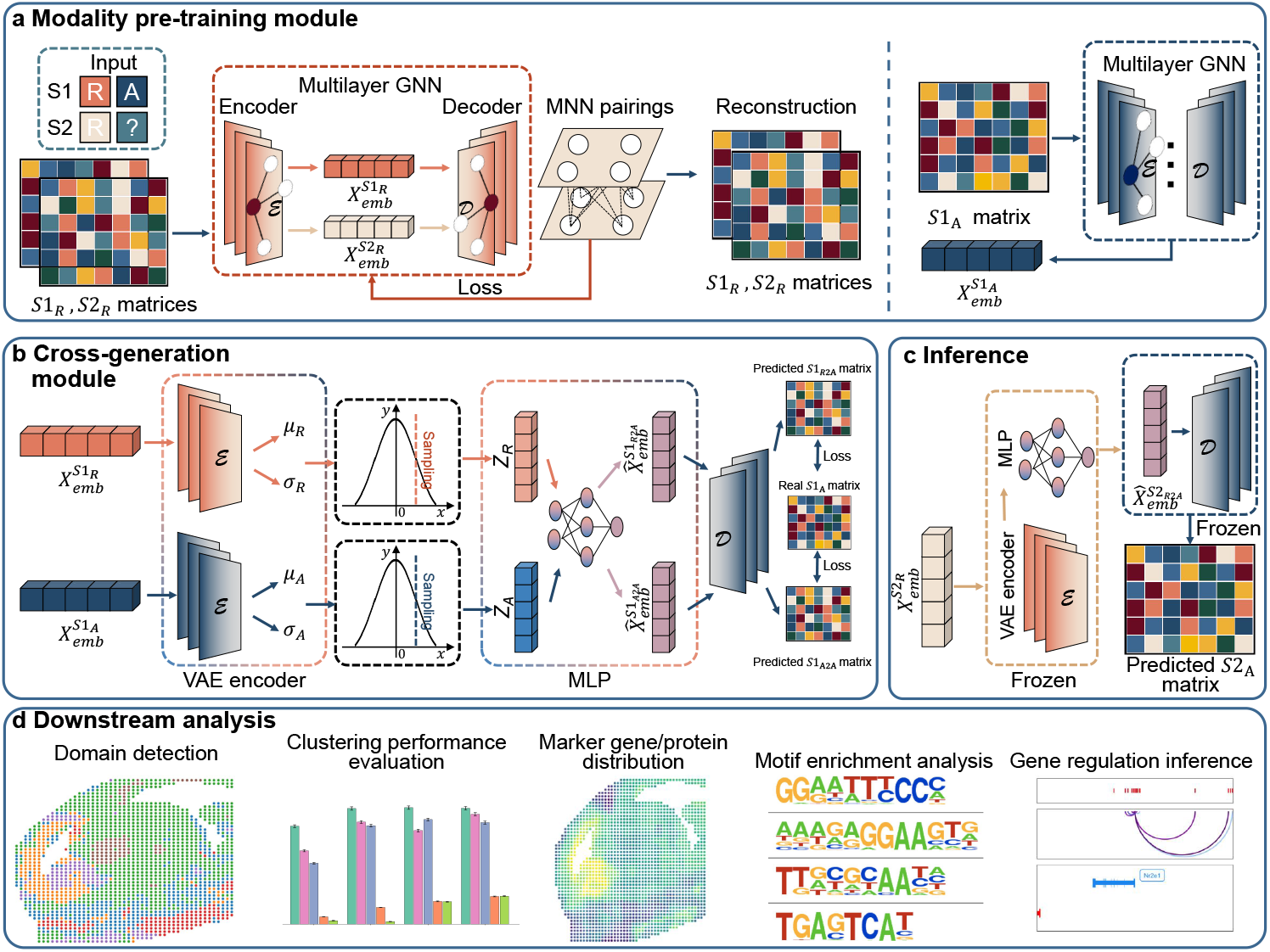
Overview of the SpaTranslator framework. We illustrate the translation from spatial RNA-seq data to spatial ATAC-seq data. Given one slice with paired spatial RNA-seq and ATAC-seq data (denoted as *S*1_*A*_ and *S*1_*R*_, respectively) and another slice with only spatial RNA-seq data (denoted as *S*2_*R*_), SpaTranslator aims to infer the missing ATAC-seq modality for the second slice (denoted as *S*2_*A*_) through the following two modules. **a** A modality-specific pre-training module is introduced to pre-train modality-specific graph neural network (GNN)-based autoencoders, to mitigate the high-dimensional sparsity of spatial omics data, thereby generating informative latent embeddings for each spatial measurement spot. In addition to the reconstruction loss, a mutual nearest neighbor (MNN)-based contrastive learning strategy is applied to the spatial RNA-seq autoencoder to remove the batch effects between slices. **b** A cross-generation module based on an adversarial variational generative model is proposed to perform cross-modality translation by cross-aligning the latent embeddings derived from the pre-trained autoencoders. MLP, multiple layer perceptron. **c** After training, the models are frozen and used to predict the missing modality. **d** The generated data are evaluated for biological fidelity and utility via domain detection, clustering performance evaluation, visualization of marker genes and proteins distribution, motif enrichment analysis, and gene regulation inference.

First, a modality-specific pre-training module is employed to train a separate autoencoder for each modality (Fig. 1a; Methods). To mitigate the high-dimensional sparsity of spatial omics data, we adopt GNN-based autoencoders that propagate information across spatially neighboring spots. For each slice, a spatial graph is constructed based on the spot coordinates. The modality-specific autoencoder takes the spatial omics matrices (i.e., *S*1_*R*_, *S*2_*R*_, and *S*1_*A*_) and the corresponding *k*-nearest neighbor (*k*NN) graphs as input, and is optimized with a reconstruction loss to learn informative latent embeddings for each spot. To mitigate batch effects across slices, a mutual nearest neighbor (MNN)-based contrastive learning strategy is applied during the training of the spatial RNA-seq autoencoder, in addition to the reconstruction objective. This module outputs informative latent embeddings for three inputs, denoted as 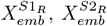, and 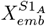 for *S*1_*R*_, *S*2_*R*_, and *S*1_*A*_, respectively, which are subsequently used by downstream modules.

Following pre-training, SpaTranslator introduces a cross-generation module based on an adversarial variational generative model, to perform cross-modality translation by cross-aligning the latent embeddings derived from the pre-trained autoencoders (Fig. 1b; Methods). Specifically, using a variational autoencoder (VAE), the embeddings 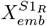 are trained to mimic the embeddings 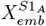, thus enabling the prediction of the ATAC-seq matrix. The VAE is optimized with a vanilla VAE loss, and an adversarial loss is further introduced during VAE training to distinguish real embeddings from generated ones, thereby further enhancing the generation quality.

After model training, all components are frozen to conduct inference for the missing modality (Fig. 1c; Methods). The RNA-seq embeddings for the target slice (i.e.,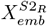) are passed to the VAE in the cross-generation module to produce the predicted spatial ATAC-seq matrix for the target slice. The generated missing modality data enables various downstream biological analyses to evaluate the reliability and biological fidelity of the predictions, including domain detection, clustering performance evaluation, spatial visualization of marker genes/proteins, motif enrichment for epigenomic data, and inference of gene regulations (Fig. 1d; Methods).

### 2.2 SpaTranslator improves intra-slice cross-modality generation performance for spatial transcriptomics and spatial epigenomics data

As a proof-of-concept, we first evaluated SpaTranslator’s performance in intra-dataset cross-modality generation using datasets obtained from MISAR-seq [4], spatial ATAC-RNA-seq and spatial CUT&Tag-RNA-seq [3] technologies, which offer spatial dual-omics profiling of the mouse brain. The MISAR-seq dataset includes joint RNA and ATAC profiling of mouse brain slices across four developmental time points (E11.0, E13.5, E15.5, and E18.5), with two slices per time point, resulting in a total of eight slices. The spatial ATAC-RNA-seq and spatial CUT&Tag-RNA-seq datasets provide joint profiling of RNA with ATAC, H3K27me3, H3K4me3 and H3K27ac for four mouse brain slices, respectively. In total, 12 slices were included in this evaluation (details in Supplementary Table 1).

The spots of each slice were randomly partitioned into training, validation, and test sets in a ratio of 7:1:2. For evaluation, one modality in the test set was removed and treated as the generation target. Following best practice [15], we performed clustering and region annotation on the generated modality from the test set and compared the resulting annotations to the original annotations using several quantitative metrics, including adjusted Rand index (ARI), normalized mutual information (NMI), adjusted mutual information (AMI), and homogeneity (HOM). To our knowledge, no previous methods have been specifically developed for cross-modality generation of spatial omics. Therefore, for performance benchmarking, we selected four representative single-cell cross-modality generation methods as baselines, including multiDGD [9], scPair [10], JAMIE [11], and scButterfly [12]. To ensure a robust evaluation, we employed five fold cross-validation and took average of five fold results to produce the final metrics.

For the average performance across the eight slices from the MISAR-seq dataset [4], SpaTranslator significantly outperformed all baseline methods across all evaluation metrics for both RNA to ATAC and ATAC to RNA translation tasks (Fig. 2a,b). Specifically, under the ATAC to RNA translation setting (Fig. 2a, left panel), SpaTranslator achieved relative improvements of 79.6% and 19.1% in terms of ARI and AMI, respectively, compared to the best performed baseline method (i.e., multiDGD). Under the RNA to ATAC translation setting (Fig. 2a, right panel), SpaTranslator achieved relative improvements of 51.7% and 16.5% in terms of ARI and AMI, respectively, compared to the best baseline method (i.e., multiDGD). For the per-slice performance, SpaTranslator consistently outperformed all baseline methods across all eight slices under both translation settings. For ATAC to RNA translation, SpaTranslator achieved relative improvements ranging from 56.8% to 96.8% in terms of ARI (Fig. 2b, left panel). For RNA to ATAC translation, it achieved relative improvements ranging from 27.7% to 85.8% in terms of ARI (Fig. 2b, right panel). Additionally, SpaTranslator achieved higher AMI scores on seven slices and higher NMI scores on six slices for both settings (Supplementary Fig. S1). These results highlighted SpaTranslator’s superior capability in cross-modality generation between RNA and ATAC.

**Fig. 2.**
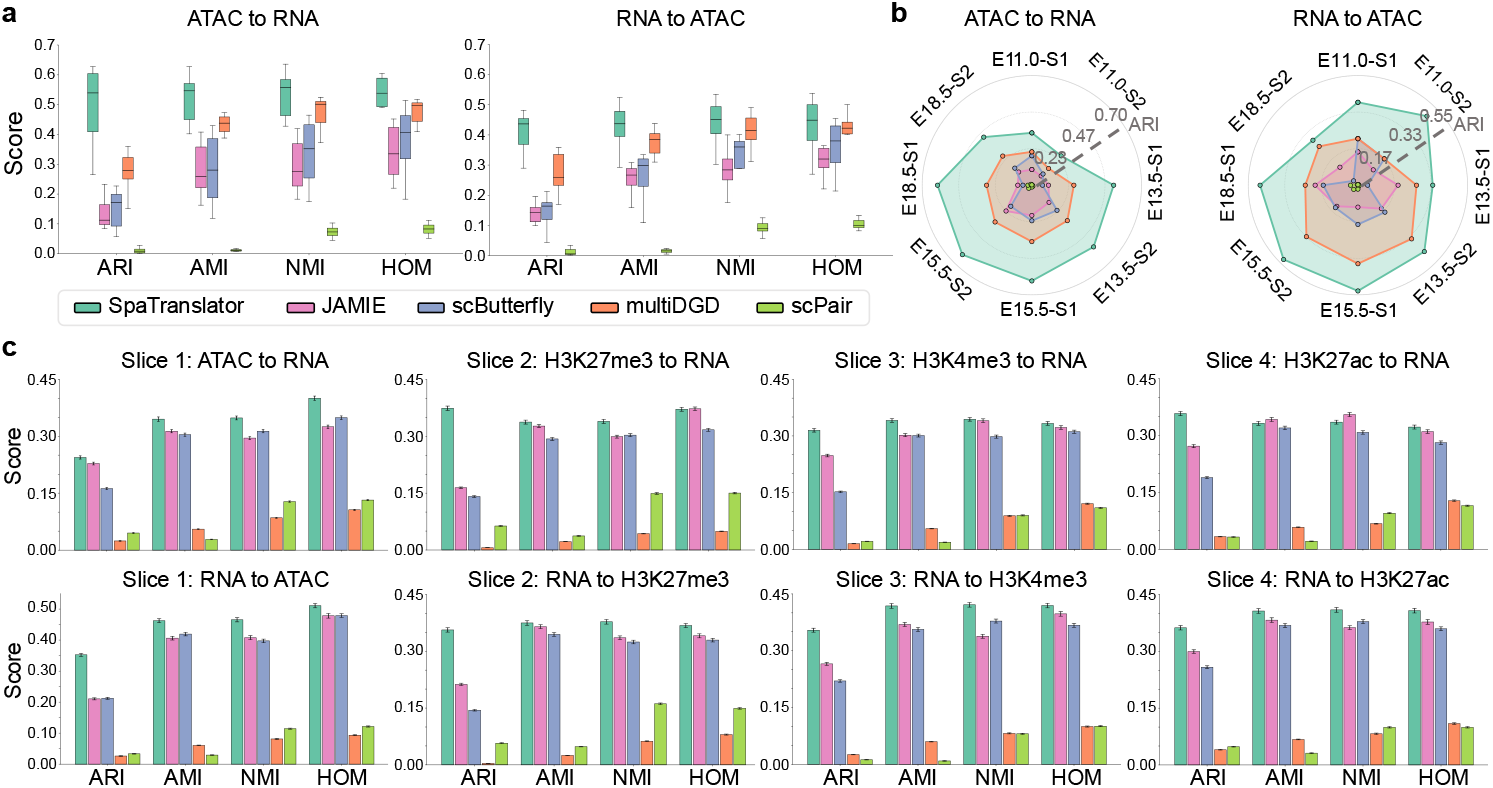
Performance evaluation on intra-slice cross-modality generation for spatial transcriptomics and epigenomics data. **a** Average clustering performance of SpaTranslator and baseline methods across eight slices from the MISAR-seq dataset, evaluated using adjusted Rand index (ARI), normalized mutual information (NMI), adjusted mutual information (AMI), and homogeneity (HOM) metrics for ATAC to RNA and RNA to ATAC translations. **b** ARI scores of SpaTranslator and baseline methods on the eight slices from the MISAR-seq dataset for ATAC to RNA and RNA to ATAC translations. E11.0, E13.5, E15.5, and E18.5 denote specific embryonic time points, while S1 and S2 represents two different slices for each time point. **c** Clustering performance on four slices from the spatial ATAC-RNA-seq and spatial CUT&Tag-RNA-seq dataset, measured using ARI, AMI, NMI, and HOM for RNA to epigenomic modalities (i.e., ATAC, H3K27me3, H3K4me3, and H3K27ac) translations and epigenomic modalities to RNA translations. All results are based on five fold cross-validation.

We further assessed SpaTranslator on the four slices from the spatial ATAC-RNA-seq and spatial CUT&Tag-RNA-seq dataset [3], which include more sequencing spots and multiple heterogeneous modality types (Supplementary Table 1). For the epigenomic to RNA translation tasks, SpaTranslator consistently outperformed all baseline methods not only in ATAC to RNA translation but also in translations from other epigenomic modalities (i.e., H3K27me3, H3K4me3, and H3K27ac) to RNA, achieving a peak relative improvement of 128.1% in terms of ARI for H3K27me3 to RNA translation (Fig. 2c, upper panel). Similarly, for the RNA to epigenomic translation tasks, SpaTranslator outperformed all baseline methods across all tasks, with a peak relative improvement of 67.6% in terms of ARI for RNA to ATAC translation (Fig. 2c, lower panel).

In summary, these results demonstrated the robustness and versatility of SpaTranslator in modality translation across diverse omics types, highlighting its capability to generate not only ATAC-seq data but also a broad range of histone modification modalities.

### 2.3 SpaTranslator enhances cross-slice cross-modality translation between spatial transcriptomics and epigenomics, yielding meaningful biological insights

Next, we evaluated SpaTranslator’s performance on cross-slice generation using two E15.5 embryonic mouse brain tissue slices from the MISAR-seq dataset [4], sourced from the left (S1) and right (S2) brains, respectively. Each slice provides joint profiling of RNA and ATAC. In our test, one modality in the right brain slice (S2) was sequentially removed and treated as the generation target (Fig. 3a). The reliability of the generated data was validated through multiple analyses: (1) comparative clustering analysis against baseline methods, including spatial visualization and quantitative metric evaluation; (2) assessment of the spatial distribution of specific biological markers, comparing marker gene expression in predicted spatial RNA data and enriched motif patterns in predicted spatial ATAC data against their respective ground truths; and (3) multi-omics integrative analysis, such as gene regulation inference, to provide novel views for explaining embryo development process.

**Fig. 3.**
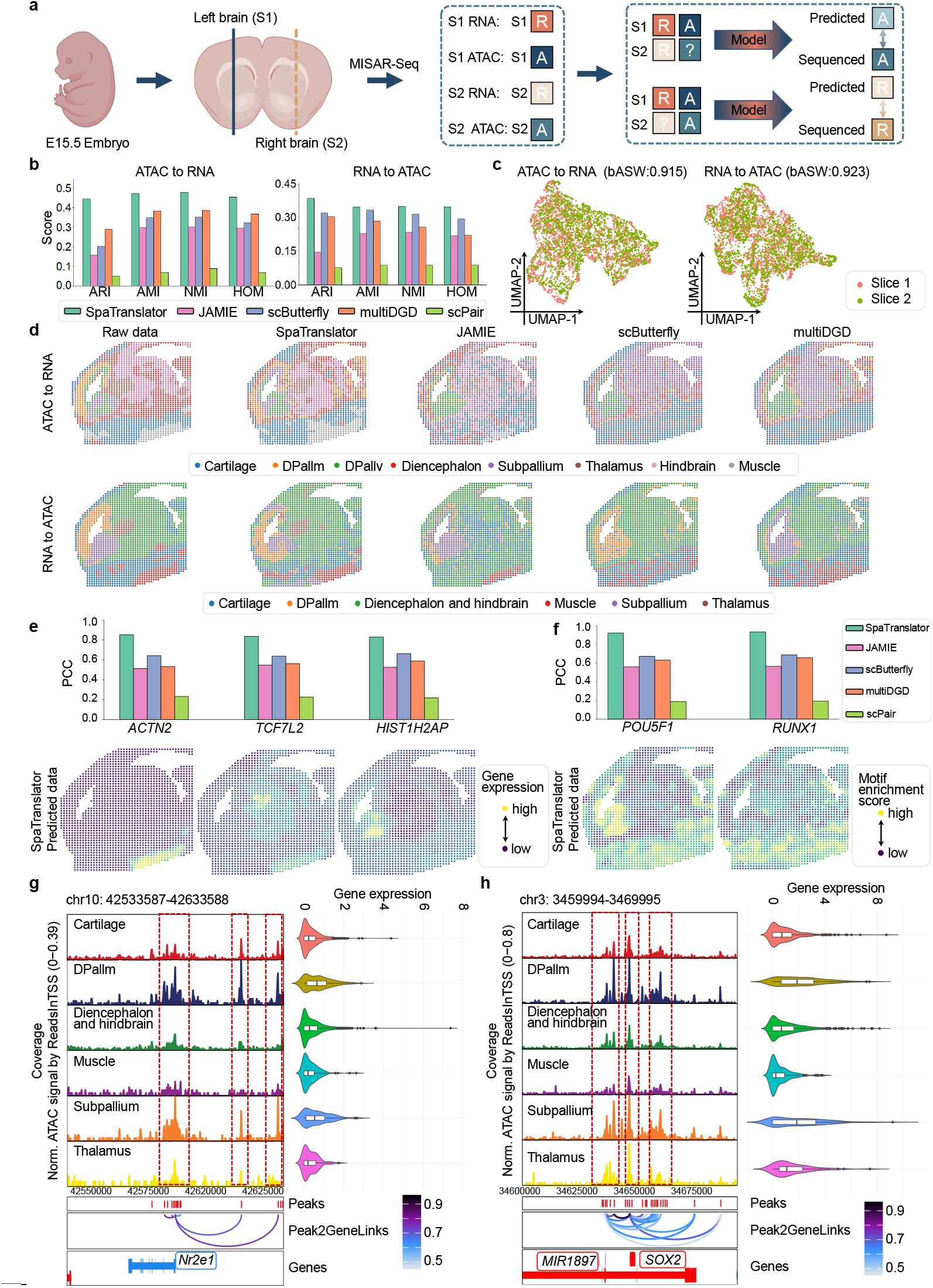
Experiment results on cross-slice cross-modality translation between spatial transcriptomics and epigenomics data. **a** Overview of the experimental workflow. Two slices from the MISAR-seq dataset were used, specifically the left brain of an E15.5 Embryo (denoted as S1) and the right brain of an E15.5 Embryo (denoted as S2), both co-profiled for RNA and ATAC modalities. For each modality in S2 (i.e., RNA or ATAC) were sequentially removed and treated as the generation target. SpaTranslator and baseline methods were then employed to predict the missing modality. **b** Clustering performance of SpaTranslator and baseline methods measured by adjusted Rand index (ARI), normalized mutual information (NMI), adjusted mutual information (AMI), and homogeneity (HOM) for ATAC to RNA and RNA to ATAC translation tasks. **c** UMAP visualizations of latent embeddings generated by SpaTranslator’s batch-correction autoencoder for cross-modality generation. Batch-correction performance was quantified using the modified average silhouette width for batch (bASW). **d** Clustering results for the raw data and predicted data obtained from SpaTranslator and baseline methods. The ground truth labels for both modalities were derived directly from the original dataset. **e** Spatial distribution maps of marker genes *ACTN2, TCF7L2*, and *HIST1H2AP*. Pearson correlation coefficients (PCCs) between predictions and ground truth are provided for SpaTranslator and baseline methods. **f** Spatial motif enrichment maps for *POU5F1* and *RUNX1*, with PCCs provided for SpaTranslator and baseline methods. **g**,**h** Regulatory maps of (**g**) *NR2E1* and (**h**) *SOX2*. Each map is organized into three panels. The upper left panel shows chromatin accessibility within a specific region (chr10:42533587–42633588 for *NR2E1*; chr3:3459994-3469995 for *SOX2*) across different domains based on SpaTranslator’s predicted ATAC data. The upper right panel displays gene expression across all domains based on actual RNA data. The lower panel depicts the genes and peaks within this genomic interval and the inferred regulatory links between them, where color intensity indicates the strength of each regulatory interaction, derived from integrated RNA–ATAC analysis. Red dashed boxes highlight potential regulatory sites.

As shown in Fig. 3b, SpaTranslator consistently outperformed all baseline methods across all metrics for the translation between RNA and ATAC. Notably, SpaTranslator achieved relative improvements of 53.1% for ATAC to RNA translation and 19.9% for RNA to ATAC translation, respectively, in terms of ARI. Moreover, in the ATAC to RNA translation task, SpaTranslator correctly identified the distinct yet encompassing relationship of the DPallv and subpallium regions, precisely delineated the muscle region, and uniquely resolved the rare Thalamus domain (Fig. 3d). In contrast, other methods struggled to separate DPallv and subpallium, producing noisy and indistinct brain region delineations. Similar fidelity was observed in the RNA to ATAC translation task, where SpaTranslator’s inferred spatial patterns closely matched the ground truth. Furthermore, visualizing the latent embeddings derived from the SpaTranslator’s batch-correction autoencoder of modality pre-training module revealed that the data from different slices exhibited highly similar manifold distributions. We then quantified the result of batch-correction by calculating the modified average silhouette width of batch (bASW). Both qualitative and quantitative results demonstrated the effective mitigation of batch effects by SpaTranslator, which contributed to its superior performance and underscored the significance of batch effects correction in the cross-modality translation for spatial omics (Fig. 3c).

We conducted ablation experiments to assess the contribution of the core design components of SpaTranslator, including the GNN and the batch-correction loss. Specifically, each component was removed in turn and the ATAC to RNA translation task was re-run on the MISAR-seq dataset. As shown in Supplementary Fig. S2a, the full model achieved the best clustering performance. Removing either the GNN or the batch-correction loss resulted in a moderate decrease in performance, underscoring their importance in mitigating data sparsity and preserving spatial characteristics. We further assessed the sensitivity of SpaTranslator to the hyperparameter *k* used in the *k*-nearest neighbor (*k*NN) graph construction. As shown in Supplementary Fig. S2b, the model’s performance remained stable across different values of *k*, demonstrating the robustness of SpaTranslator to this hyperparameter.

To further validate the reliability of the spatial data generated by SpaTranslator, we analyzed the spatial distribution of marker genes and motifs using spatial distribution maps (Fig. 3e,f). *ACTN2* is known for its specific expression and critical roles in the muscle region of the mouse brain [16–18]. As shown in the gene expression map (Fig. 3e), *ACTN2* displayed high, localized expression in the muscle region, with sharp boundaries separating it from other brain areas. This trend was further supported by other marker genes. For instance, *TCF7L2* a transcription factor essential for thalamus development and functional maintenance, regulates the differentiation, maturation, and gene expression of dorsal thalamic neurons [19]. In SpaTranslator’s predicted spatial expression map, *TCF7L2* showed enriched expression specifically within the thalamus region. Similarly, *HIST1H2AP*, predominantly expressed in a proliferative microglial subpopulation that contributes to neural stem cell proliferation and differentiation [20], was observed to be enriched in the subpallium region in the generated data. We further quantified prediction accuracy by calculating the Pearson correlation coefficient (PCC) between the predicted and real expression values for these marker genes. SpaTranslator consistently achieved PCC scores above 0.8, with the highest relative improvement reaching up to 37.1% compared to baseline methods, thereby reinforcing the reliability of its generated spatial data (Fig. 3e).

Due to the high sparsity of peak signals, we applied an enrichment analysis workflow to validate the reliability of the spatial ATAC data generated by SpaTranslator. First, we identified highly variable peaks and then performed motif enrichment analysis using HOMER [21]. The enriched motifs were subsequently matched to the JASPAR database [22] to identify biologically meaningful transcription factor motifs. The identified motifs were visualized in spatial distribution maps (Fig. 3f), enabling the identification of region-specific motifs within their spatial context. Notably, the *POU5F1* motif was highly enriched in the subpallium region, consistent with previous reports describing its role in co-binding specific enhancer sequences to activate downstream gene expression and regulate neural stem cell proliferation and differentiation during embryonic development [23]. Similarly, the *RUNX1* motif was enriched in the muscle region, in agreement with previous findings showing its involvement in myogenic differentiation and in signal transduction in muscle cells [24]. We further quantified the similarity between generated and real data by computing the PCC scores between their motif enrichment scores. SpaTranslator outperformed baseline methods achieving PCC scores higher than 0.8, whereas most baseline methods scored below 0.7 (Fig. 3f).

Next, we explored whether generating additional modalities from unimodal data could enhance multi-omics integrative analysis and offer novel biological insights. Specifically, we first used SpaTranslator to predict spatial ATAC data from the corresponding real spatial RNA profiles. Leveraging both modalities, we then performed integrative analysis using the ArchR toolkit [25] to compute peak-to-gene associations and visualize regulatory links, thereby revealing domain–specific regulatory relationships. We presented two representative cases: *NR2E1* and *SOX2* (Fig. 3g,h). The *NR2E1* gene is a well-established regulator of neural stem cell proliferation and differentiation [26]. Previous studies report that the DPallm brain region is enriched in neural stem cells that differentiate into various neuronal subtypes, while the subpallium serves as a primary source of the basal ganglia [27]. In our analysis, *NR2E1* showed relatively higher expression in the DPallm and subpallium regions compared to other brain regions. These two regions also exhibit correspondingly higher chromatin accessibility at its promoter region, indicating that the predicted ATAC data and the actual RNA data were consistently aligned in reflecting the gene’s biological significance (Fig. 3g). Moreover, integrated analysis results further revealed two distal regulatory elements associated with *NR2E1*, both displaying domain-specific high accessibility in DPallm and subpallium. Similarly, *SOX2*, another key developmental gene in the brain, showed consistent and biologically plausible regulatory patterns in our multi-omics analysis (Fig. 3h). Collectively, these results demonstrate that SpaTranslator enables effective integrative multi-omics analysis even when only unimodal data are available, overcoming the critical limitation of single-modality datasets in gene regulation studies and providing novel views into regulatory mechanisms and developmental processes.

In summary, comprehensive evaluations, including clustering, marker gene visualization, motif enrichment analysis, and gene regulation inference, collectively validated the reliability of the data generated by SpaTranslator and demonstrated its superior ability to reveal biological insights into mouse brain developmental processes.

### 2.4 SpaTranslator enables effective cross-slice, cross-modality generation for spatial transcriptomics and proteomics, allowing the revelation of immune hotspot microenvironments

After validating SpaTranslator’s superior capability in the cross-modality translation between spatial epigenomics and transcriptomics, we extended its application to the cross-modality generation between spatial proteomics and transcriptomics, to further assess its generalizability and biological relevance across a broader range of spatial omics contexts.

In this test, we employed a human tonsil dataset and a human lymph node dataset generated by the 10x Genomics Visium spatial RNA and protein co-profiling platform [28]. The tonsil is a complex immune organ composed of B cell follicles embedded in T cell enriched zones, surrounded by epithelial and connective tissues. Within the follicles, germinal centers transiently form to support B cell activation and proliferation during immune responses [29]. The lymph node is another critical immune organ, featuring a layered structure of cortex and medulla, encased by a fibrous capsule [30]. These datasets provided complex spatial microenvironment and biological scenarios, offering a robust setting to demonstrate SpaTranslator’s capability.

First, we evaluated SpaTranslator using two tissue slices from the human tonsil dataset, each providing joint profiling of RNA and protein, with protein quantified via antibody-derived tags (ADTs). In our experiment, one slice was designated as the target slice for generation, and one modality in this slice was sequentially removed and treated as the generation target. We conducted clustering analyses and plotted spatial distribution maps using the generated data. SpaTranslator consistently outperformed all baseline methods across all metrics for both ADT to RNA and RNA to ADT translation tasks (Fig. 4a). Moreover, SpaTranslator produced the most coherent regional assignments, closely aligning with ground truth annotations (Fig. 4c).

**Fig. 4.**
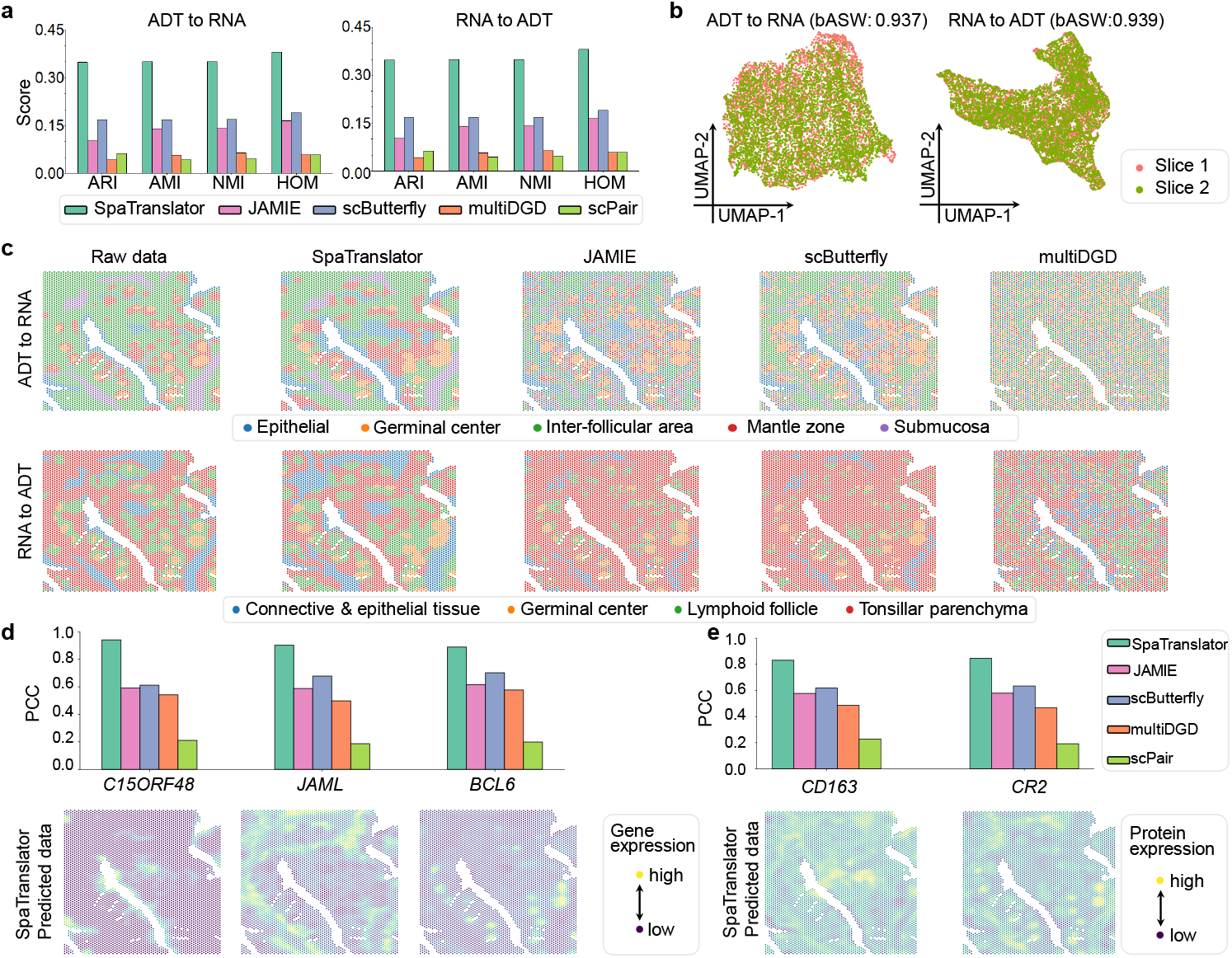
Experiment results on cross-slice cross-modality generation for spatial transcriptomics and proteomics data on human tonsil dataset. **a** Clustering performance of SpaTranslator and baseline methods, measured by adjusted Rand index (ARI), normalized mutual information (NMI), adjusted mutual information (AMI), and homogeneity (HOM) for ADT to RNA and RNA to ADT translation tasks. **b** UMAP visualizations of latent embeddings derived from SpaTranslator’s batch-correction autoencoder for cross-modality generation. Batch-correction performance was quantified using the modified average silhouette width for batch (bASW). **c** Clustering results for the raw data and predicted data obtained from SpaTranslator and baseline methods. The ground truth labels for both modalities were derived directly from the original dataset. **d** Spatial distribution maps of marker genes *C15ORF48, JAML*, and *BCL6*. Pearson correlation coefficients (PCCs) between predictions and ground truth are provided for SpaTranslator and baseline methods. **e** Spatial distribution maps of marker proteins CD163 and CR2, with PCCs provided for SpaTranslator and baseline methods.

In the ADT to RNA task, SpaTranslator accurately recovered the spatial localization of the germinal center and mantle zone. Consistent with biological knowledge [29], the mantle zone enveloped the germinal center, the latter being a key site for B cell proliferation and differentiation. The model also faithfully reconstructed the inter-follicular area, epithelial, and submucosa regions. In contrast, baseline methods such as JAMIE and scButterfly identified only the germinal center, producing fragmented and noisy representations of other regions, while MultiDGD and scPair failed to recover any meaningful structure. Additionally, as observed previously, SpaTranslator effectively mitigated batch effects when integrating same-modality data from different slices, yielding comparable latent embedding distributions across batches (Fig. 4b).

SpaTranslator also demonstrated highly faithful reconstructions of the spatial distributions of key marker genes (Fig. 4d). *C15ORF48*, an epithelial-specific gene that modulates mitochondrial metabolism and inflammatory responses under stimulation [31], showed elongated expression patterns in epithelial areas in SpaTranslator’s predictions. Similarly, *BCL6*, essential for germinal center formation and B cell maturation [32], exhibited enriched expression within follicular regions, while *JAML*, involved in T cell activation and migration [33], was more enriched in the inter-follicular area. To quantitatively assess prediction accuracy, we calculated PCC scores between the predicted and real spatial expression values of these marker genes. The predictions from SpaTranslator closely matched the ground truth, with PCC scores exceeding 0.9 for *C15ORF48* and *JAML*, indicating strong positive correlations. In contrast, baseline methods failed to achieve a comparable level of spatial reconstruction fidelity.

For the protein data generated by SpaTranslator, favorable and biologically consistent spatial patterns of key marker proteins were also observed (Fig. 4e). For instance, CR2, which synergizes with B cell receptors to enhance antigen presentation [34], showed high expression in the germinal centers, while CD163, an immunoregulatory marker expressed on macrophages [35], was localized to the inter-follicular area. Quantitative evaluation (PCC) further confirmed the reliability of these predictions.

We further evaluated SpaTranslator on the human lymph node dataset to assess its generation performance (Fig. 5). SpaTranslator again achieved superior clustering performance compared to all baseline methods (Fig. 5a). In the ADT to RNA translation task, SpaTranslator accurately delineated major lymph node structures, including the outer adipose tissue and capsule, the paracortex layer, and the internal medullary cords and sinus. In contrast, baseline methods either failed to detect these regions (i.e., multiDGD and scPair) or lacked spatial accuracy (i.e., JAMIE and scButterfly), particularly mislocalizing the capsule and adipose tissue (Fig. 5c). In the RNA to ADT translation task, SpaTranslator not only reconstructed these major domains but also uniquely identified rare regions such as the hilum and follicle. The hilum, the entry and exit point for lymphatic and blood vessels, is crucial for anatomical identification, while the follicle serves as a key immune activation site [30]. However, none of the baseline methods were able to reconstruct these structures, instead producing noisy and incoherent clustering results (Fig. 5c). Furthermore, SpaTranslator effectively mitigated batch effects when integrating same-modality data from different slices, yielding highly comparable latent embedding distributions across batches (Fig. 5b).

**Fig. 5.**
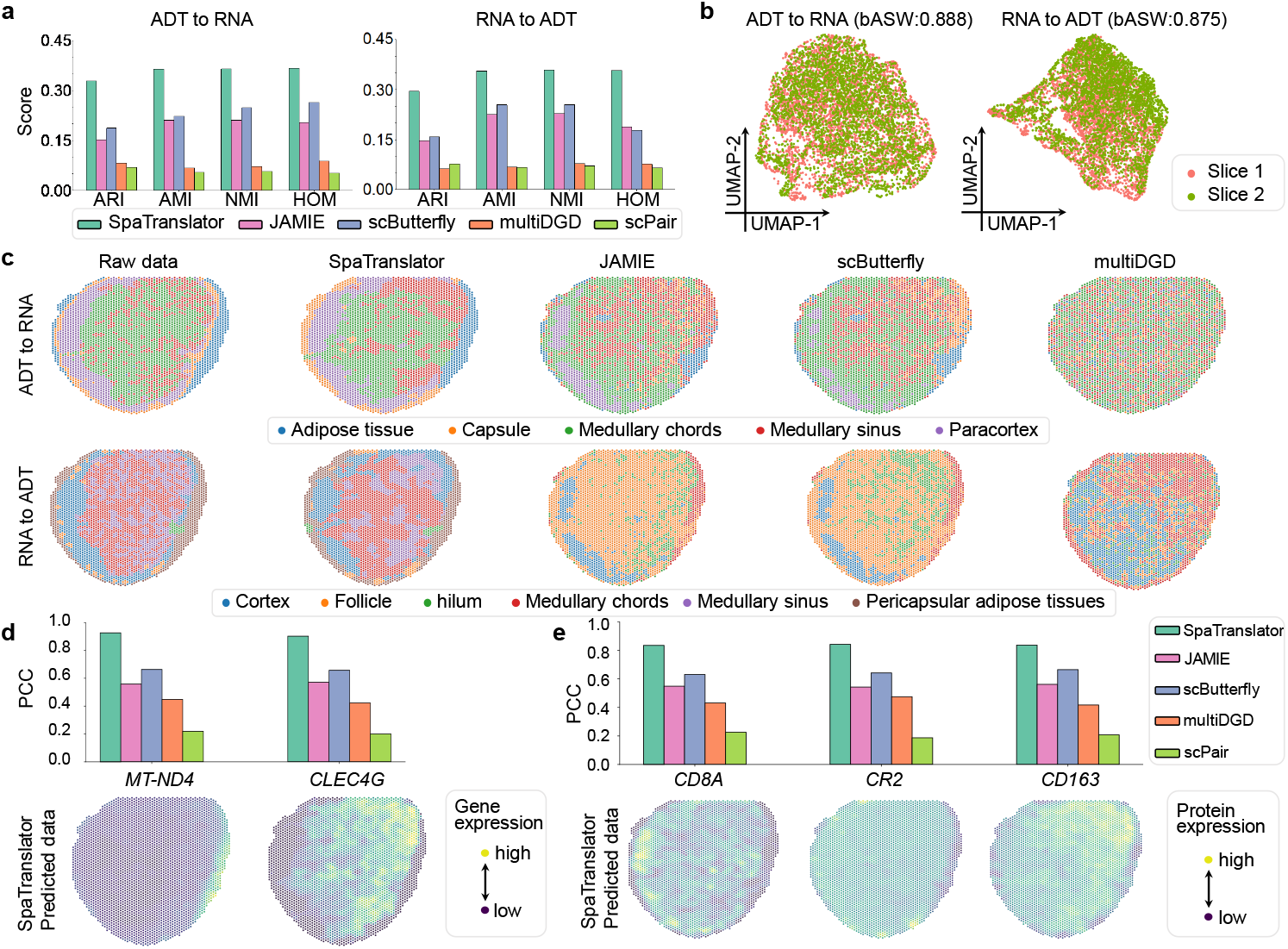
Experiment results on cross-slice, cross-modality generation for spatial transcriptomics and proteomics data on human lymph node dataset. **a** Clustering performance of SpaTranslator and baseline methods measured by adjusted Rand index (ARI), normalized mutual information (NMI), adjusted mutual information (AMI), and homogeneity (HOM) for ADT to RNA and RNA to ADT translation tasks. **b** UMAP visualizations of latent embeddings derived from SpaTranslator’s batch-correction autoencoder for cross-modality generation. Batch-correction performance was quantified using the modified average silhouette width for batch (bASW). **c** Clustering results for the raw data and predicted data obtained from SpaTranslator and baseline methods. The ground truth labels for both modalities were derived directly from the original dataset. **d** Spatial distribution maps of marker genes *MT-ND4* and *CLEC4G*, with pearson correlation coefficients (PCCs) of SpaTranslator and baseline methods are provided. **e** Spatial distribution maps of marker proteins CD8A, CR2, and CD163, with PCCs provided for SpaTranslator and baseline methods.

We then validated the biological fidelity of SpaTranslator’s predictions by examining the spatial expression of marker genes and proteins in the lymph node (Fig. 5d,e). *MT-ND4*, highly expressed in adipose tissue and involved in metabolic regulation and inflammation [36], showed peripheral localization. *CLEC4G*, a C-type lectin expressed in the medulla and involved in antigen presentation [37], was accurately mapped to the medullary region. At the protein level, CD8A was localized to the cortex, consistent with its role in T cell activation [38]. As in the tonsil dataset, CR2 and CD163 were again accurately reconstructed, further validating that the predicted data patterns align with existing literature and hold biological significance. CR2 is expressed in follicular dendritic cells and B cells, playing a central role in immune complex trafficking and B cell activation [39], while CD163 is a hallmark of macrophages and monocytes, enriched in the medulla and involved in antigen processing [40]. Quantitative analysis further confirmed a high degree of similarity between the predicted data and the ground truth, surpassing the performance of all baseline methods.

In summary, SpaTranslator performed accurate cross-modality generation between spatial transcriptomics and proteomics, enabling the revelation of immune hotspot microenvironments. Moreover, CD163 and CR2 were accurately reconstructed in both the human tonsil and lymph node datasets, demonstrating the robustness and generality of our model in reconstructing key immune markers across various organs.

## 3 Discussion

In this study, we present SpaTranslator, a deep generative framework designed for universal cross-modality generation of spatial multi-omics data. By integrating a GNN-based modality pre-training module with an adversarial generative model-based cross-generation module, SpaTranslator effectively addresses the sparsity and noise inherent in spatial omics data, mitigates batch effects, and enables accurate, robust cross-modality translation.

Through extensive evaluations on various spatial multi-omics datasets across different modalities, species, and experimental conditions, SpaTranslator consistently outperformed all baseline methods, significantly enhancing the accuracy and reliability of both inter-slice and cross-slice translations of multi-modal spatial omics data. UMAP visualizations of latent embeddings obtained from its batch-correction autoencoder of the modality pre-training module further demonstrated effective removal of batch effects across slices, contributing to its superior performance. Moreover, analyses of the spatial distributions of marker genes and proteins, together with motif enrichment assessments, confirmed that SpaTranslator faithfully reproduces the biological patterns observed in real data. Moreover, by generating missing modalities from unimodal spatial omics data, SpaTranslator enables integrative multi-omics investigations of gene regulatory relationships, offering novel perspectives for interpreting the developmental processes of the mouse brain. In summary, SpaTranslator is a powerful and versatile framework for cross-modality translation of spatial omics.

There are several avenues for future improvement and expansion of SpaTranslator. First, with the growing clinical adoption of imaging-based omics such as histology, computed tomography, and magnetic resonance imaging, incorporating image data as an input modality could further enhance generative accuracy and reduce reliance on costly spatial sequencing, enabling more cost-effective and fine-grained investigations into disease mechanisms. Second, while SpaTranslator currently supports spatial dual-omics inputs, emerging experimental techniques such as spatial-Mux-seq [41] enable simultaneous spatial profiling of more than two modalities. Extending our framework to handle multi-modal inputs would allow for mosaic generation—inferring missing modalities from partial inputs, thereby further broadening the model’s utility. Finally, SpaTranslator is designed to model two-dimensional tissue slices. Exploring extensions to three-dimensional (3D) tissue reconstruction would open new avenues for constructing virtual organs and simulating complex biological processes, such as tumorigenesis and organ development in a fully 3D spatial context.

## 4 Methods

### 4.1 Data collection and preprocessing

To comprehensively evaluate the performance of the SpaTranslator model, we collected multiple spatial multi-omics datasets (Supplementary Table 1). For the spatial transcriptome-epigenome translation task, we used two datasets generated by MISAR-seq [4], spatial ATAC-RNA-seq and spatial CUT&Tag-RNA-seq [3] technologies. All of the technologies measure mouse embryonic brain tissue and enable spatial dual-omics profiling of the transcriptome and epigenome. The MISAR-seq dataset includes four developmental time points of the mouse brain (i.e., E11.0, E13.5, E15.5, and E18.5), with two slices measured at each time point (one left brain and one right brain). The spatial ATAC-RNA-seq and spatial CUT&Tag-RNA-seq dataset provides transcriptomics and epigenomics co-profiling for four slices, including one with RNA and ATAC co-profiling, and three slices for RNA and three histone modifications (i.e., H3K27me3, H3K4me3, and H3K27ac), respectively. Collectively, these datasets provided 12 distinct slices for the spatial transcriptome-epigenome cross-modality translation task.

For the spatial proteome-transcriptome cross-modality translation task, we utilized two datasets generated by the 10x Genomics spatial Visium RNA and protein co-profiling platform [28]: a human tonsil dataset and a human lymph node dataset. Both datasets originate from immune-related tissues and provide insight into the human immune microenvironment.

We applied distinct preprocessing methods tailored to different modalities. For the spatial transcriptomics count matrix, we normalized the total count of each cell to match the median total counts across all cells prior to normalization. The normalized values were subsequently log-transformed with an offset of 1. We then selected the top 3,000 highly variable genes (HVGs) to serve as input to the model [15].

For spatial ATAC-seq data, we first binarized the matrix and filtered out peaks that were activated in fewer than 0.5% of cells. We then applied a term frequency-inverse document frequency (TF-IDF) transformation [42]. Following this, the matrix was scaled to the range [0,1].

For the spatial proteomics count matrix, we performed the centered log ratio (CLR) transformation across cells, as defined by:

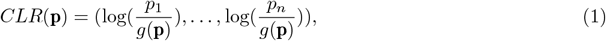

where **p** = (*p*_1_, …, *p*_*n*_) represents the count vector of protein epitopes for each cell, and *g*(**p**) denotes the geometric mean of vector **p**.

In addition to the count matrices, SpaTranslator requires a spatial adjacency graph as input. We constructed a *k*NN graph based on the spatial coordinates of the spots, with each spot represented as a node, and edges derived from the Euclidean distances between the spots [43]. We set *k* = 6 in our experiments, meaning that the six nearest spots were considered as neighbors for each spot. This yielded an undirected adjacency matrix *A*, where *A*_*ij*_ = 1 if spot *i* is among the *k* nearest neighbors of spot *j*, and 0 otherwise. The matrix *A* is symmetric and includes self-loops, i.e., *A*_*ii*_ = 1.

### 4.2 SpaTranslator model details

To illustrate the framework of SpaTranslator, consider the task of translating spatial RNA-seq data into spatial ATAC-seq data. Given a slice (denoted as S1) with paired spatial RNA-seq (denoted as *S*1_*R*_) and ATAC-seq (denoted as *S*1_*A*_) data, and another slice (denoted as S2) with only spatial RNA-seq data (denoted as *S*2_*R*_), SpaTranslator aims to leverage the paired data from the S1 as a reference to infer the missing ATAC-seq modality (denoted as *S*2_*A*_) for S2. SpaTranslator consists of two main modules: a modality pre-training module and a cross-generation module.

#### 4.2.1 Modality pre-training Module

A modality-specific pre-training module is introduced to pre-train individual autoencoders for RNA-seq and ATAC-seq modalities, respectively (Fig. 1a). To enable effective feature representation and address the high-dimensional sparsity characteristics of spatial omics data, GNN-based autoencoders are employed to enable the interaction of information among neighboring spatial locations.

For the RNA-seq modality, the autoencoder focuses on generating informative feature representations while simultaneously correcting batch effects across slices. Specifically, suppose we have two RNA-seq matrices *S*1_*R*_ and *S*2_*R*_, along with their corresponding *k*NN graph 𝒢_*S*1_ and 𝒢_*S*2_ for S1 and S2, respectively. A graph attention network (GAT)-based encoder [44] (denoted as Encoder_*R*_) takes these matrices and graphs as inputs to generate latent representations as follows:

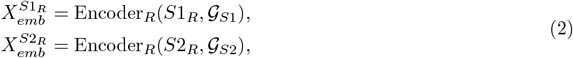

where 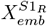 and 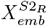 denote the latent embeddings for *S*1_*R*_ and *S*2_*R*_, respectively. Then a GNN-based decoder (denoted as Decoder_*R*_) reconstructs the inputs from the latent embeddings as follows:

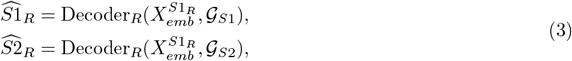

where 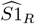 and 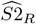 denote the reconstructed matrices. The reconstruction loss is defined as the mean squared error (MSE) between the reconstructed matrices and their original inputs:

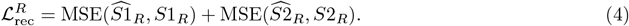

To eliminate inter-sample batch effects between slices, we adopt an MNN-based contrastive learning strategy, guiding the encoder to produce batch-invariant embeddings. This strategy relies on triplet-based learning in the latent space. Each triplet consists of an anchor, a positive, and a negative spot. The anchor and positive spots are MNN-paired spots from different slices with similar gene expression profiles, while the negative is randomly selected from the same slice as the anchor. A contrastive loss is employed to encourage the anchor to be closer to the positive while pushing it away from the negative:

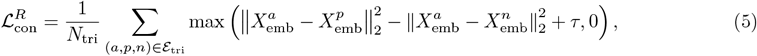

where 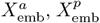, and 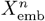 denote the anchor, positive, and negative spot embeddings derived from 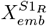 or 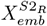, respectively. ℰ_*tri*_ denotes the set of triplets. *τ* denotes a margin hyperparameter and is set to 1 by default.

The total loss for the RNA-seq autoencoder combines the reconstruction loss and contrastive loss as follows:

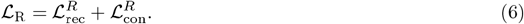

For the ATAC-seq modality, the autoencoder is trained to generate informative feature representations for individual spots and to accurately reconstruct ATAC-seq profiles, thereby enabling the generation of missing ATAC-seq data. We adopt a VAE framework [45]. Specifically, given the ATAC-seq matrix *S*1_*A*_ and the corresponding *k*NN graph 𝒢_*S*1_, a graph convolutional network (GCN)-based encoder (denoted as Encoder_*A*_) takes these as inputs and outputs the latent embeddings 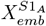:

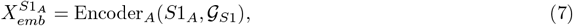

Subsequently, two multilayer perceptrons (MLPs) are employed to map 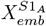 to a mean vector *µ*_*A*_ and log-variance vector *σ*_*A*_, respectively. Using the mean and log-variance, the reparameterization trick is used to sample from the latent distribution, resulting in the sampled embedding *Z*_*A*_ as follows:

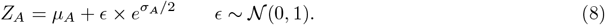

An MLP *T* (·) further transforms *Z*_*A*_ into intermediate latent embeddings 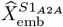, which are then used as input to an MLP-based decoder to reconstruct the original ATAC-seq matrix as follows:

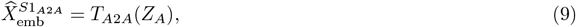

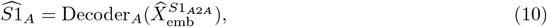

The reconstruction loss is computed using the binary cross-entropy (BCE) between the reconstructed and original matrices as follows:

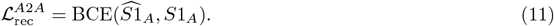

The total loss for the ATAC-seq variational autoencoder combines the reconstruction loss and evidence lower bound (ELBO) loss, which regularizes the latent distribution towards a standard normal prior:

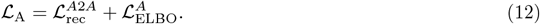

#### 4.2.2 Cross-generation module

After pre-training, SpaTranslator incorporates a cross-generation module based on an adversarial variational generative model to achieve cross-modality generation. The latent embeddings for RNA-seq matrix derived from the modality pre-training module (i.e., 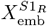), are fed into an MLP-based encoders to derive mean and log-variance vectors (denoted as *µ*_*R*_ and *σ*_*R*_). Subsequently, the latent embeddings *Z*_*R*_ are sampled using the reparameterization trick. Then, an MLP *T*_*R*2*A*_(·) is used to perform cross-modality mapping in latent space, and the Decoder_*A*_ pre-trained in the modality pre-training module is finetuned to decode the original ATAC-seq matrix as follows:

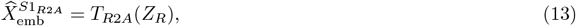

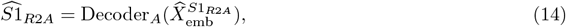

where 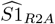 denotes the predicted ATAC-seq data translated from the RNA-seq feature embeddings. A BCE loss (denoted as 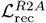) is used to measure the deviation between the 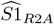 and *S*1_*A*_ and an ELBO loss (denoted as 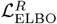) is used to regularize the latent distribution of *Z*_*R*_ towards a standard normal prior.

To make the generated embedding more similar to the original, we introduced a generative adversarial network (GAN) model for adversarial training [46]. Specifically, a discriminator *D*(·) is trained to discriminate the generated and original embeddings and the cross-generation module is trained to mimic the distribution of the original embedding. When the discriminator cannot distinguish between real and fake, the model training has achieved the desired effect. To compute the discriminator’s loss, we implemented the soft labels technique. Specifically, we first sample the smoothed positive label *l*_*pos*_ ∼ Uniform[0.8, 1] and negative label *l*_*neg*_ ∼ Uniform[0, 0.2], respectively for the labels 1 and 0 [47]. The loss function for the discriminator is defined as follows:

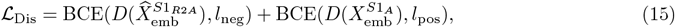

To avoid the catastrophic forgetting problem of the pre-trained Decoder_*A*_, the ATAC-seq VAE loss (i.e., ℒ_*A*_) is introduced in the training of this module. Finally, the total loss for the cross-generation module is defined as follows:

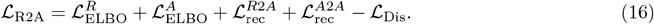

#### 4.2.3 Inference module

Following model training, all components are frozen to perform inference for the missing modality. The RNA-seq embeddings of the target slice (denoted as 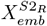) are passed to the cross-generation module to obtain the predicted ATAC-seq latent embeddings (denoted as 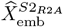) and then decoded by the ATAC decoder to produce the predicted spatial ATAC-seq matrix for the target slice.

#### 4.2.4 Model configurations

##### Modality pre-training module

The RNA-seq autoencoder comprises a two-layer GNN encoder with hidden dimensions 512 and 128 and a two-layer GNN decoder with hidden dimensions 128 and 512. The ATAC-seq autoencoder comprises a two-layer GNN encoder with hidden dimensions 512 and 128 and a two-layer MLP decoder with hidden dimensions 128 and 512. Parameters for both models were optimized using the Adam optimizer [48] with a learning rate of 0.001. The RNA model was trained for 500 epochs using only a reconstruction loss; after this warm-up, training continued for an additional 500 epochs during which a contrastive batch-correction loss was applied every 10 epochs. The ATAC model was trained for 200 epochs using the VAE loss.

##### Cross-modality generation module

The latent embedding dimension for each spot was set to 128. Each fully connected layer employed a LeakyReLU activation function. For the discriminator, a Sigmoid output layer was appended after the fully connected layers. The model was trained for 200 epochs using the Adam optimizer with a learning rate of 0.001 for parameter optimization. An early-stopping strategy was implemented, terminating training if the loss did not significantly improve over 50 consecutive epochs.

All experiments were conducted on a server with four-core Intel(R) Xeon(R) Platinum 8358 CPUs @ 2.60GHz and a single NVIDIA A100 GPU (80G).

### 4.3 Downstream analysis

To comprehensively evaluate the model’s performance, we apply a range of analytical approaches to the predicted data, including clustering, marker gene/protein visualization, motif enrichment analysis, and gene regulation inference.

#### 4.3.1 Clustering and visualization

For predicted data with dimensionality greater than 50, principal component analysis (PCA) was first applied to reduce the dimensions to 50. For low-dimensional proteomic data, subsequent steps were applied directly without dimensionality reduction. The Leiden algorithm, implemented via the `Scanpy` package [49], was used to cluster the data and identify spatial domains.

To incorporate prior knowledge, the number of anatomical regions (e.g., brain regions) was predefined, and a grid search was used to determine the optimal resolution parameter that matched the target number of clusters. If grid search could not find optimal resolution we performed cell clustering by the Leiden algorithm with default resolution of one [50]. Then, each cluster was annotated based on its markers. After cluster labels were assigned, the `sc.pl.embedding()` function was utilized to visualize the spatial distribution of spots on a two-dimensional plane, with each spot colored by its cluster labels. By comparing clustering results from different methods alongside the ground truth on the same canvas, the qualitative accuracy of clustering was assessed.

Moreover, to assess the performance of batch-correction, we visualized latent embeddings derived from the SpaTranslator’s batch-correction autoencoder of the modality pre-training module to check whether data from different slices exhibited highly similar manifold distributions. We then quantified the result of batch-correction by calculating bASW following a previous benchmark study [51].

#### 4.3.2 Evaluation metrics

We quantitatively evaluated the clustering performance of SpaTranslator uisng four standard metrics, including ARI, AMI, NMI, and HOM. The Rand index (RI) measures the similarity between predicted and true labels, while ARI adjusts RI to account for chance agreement. Mutual information (MI) quantifies the dependence between predictions and true labels, scaled by NMI and further corrected for chance by AMI. HOM assesses cluster purity, with a score of 1 indicating that each cluster contains only a single cell type. Additional details on these evaluation metrics, are provided in Supplementary Notes C.

#### 4.3.3 Implementation details of baseline methods

We compared SpaTranslator with four representative methods for the modality translation in single cell data, including multiDGD, scPair, JAMIE, and scButterfly. Each method was executed following its publicly available GitHub instructions, using default hyperparameters. All experiments were conducted under identical hardware settings. Notably, these four baselines were originally designed for cross-modality generation in single-cell data and therefore operated solely on raw count matrices, without incorporating spatial graph structures. A detailed methodological comparison between SpaTranslator and the baseline methods is provided in Supplementary Table 2.

#### 4.3.4 Visualization of marker genes and proteins

Cluster-specific marker genes and proteins were identified using the `rank_genes_groups` function from the `Scanpy` package [49], selecting the top five highly variable features for each cluster. The spatial distributions of these features were then visualized for the predicted data. To validate biological relevance, we cross-referenced published literature to confirm that the identified genes and proteins were indeed region-specific and functionally important in those locations. To quantitatively assess the similarity between the generated and the real data, we calculated PCCs of expression values for specific genes across all spots between the generated data and the corresponding ground truth data. These results were subsequently compared with those obtained from baseline methods.

#### 4.3.5 Motif enrichment analysis

For epigenomic data, motif enrichment analysis was performed following established procedures [52]. Cluster-specific peaks were identified using the `rank_genes_groups` function with the Wilcoxon test. Peaks with a log_2_ fold change below 0.2 were excluded, and the top 1000 peaks with the smallest adjusted *p*-values (q-values) were retained for each cluster. These peaks were then compiled into a peak set for each cluster and analyzed using the `chromVAR` package (v1.26.0) [52] to identify enriched transcription factor motifs. Enrichment scores were computed and mapped to the JASPAR database [22] to identify biologically significant motifs. The spatial distributions of enriched motifs were then visualized and compared between predicted and real data. PCC was calculated to quantify the similarity between generated data and the corresponding ground truth.

#### 4.3.6 Gene regulation inference

To infer regulatory relationships, we integrated the real RNA and predicted ATAC datasets and then employed the ArchR toolkit [25] to compute peak-to-gene correlations. Significant regulatory links were identified using ArchR’s `addPeak2GeneLinks` and `getPeak2GeneLinks` functions with default settings. These links were visualized using the `plotBrowserTrack` function, which displays chromatin accessibility, gene expression levels, and the strength of peak-gene associations across different cell types, thus revealing cell type-specific regulatory architectures.

## 5 Data availability

All data used in this study are publicly available. The MISAR-seq data analyzed in this study are accessible through the National Genomics Data Center under accession number OEP003285 (https://www.biosino.org/node/project/detail/OEP003285). Raw and processed spatial ATAC-RNA-seq and spatial CUT&Tag-RNA-seq data are available in the Gene Expression Omnibus (GEO) under accession number GSE205055. The 10x Genomics Visium spatial RNA and protein co-profiling datasets from human lymph node and tonsil can be accessed at https://zenodo.org/uploads/12654113.

## 6 Code availability

All code for SpaTranslator, including model implementation and tutorials is available at: https://github.com/donghongyu2020/SpaTranslator.

## 7 Acknowledgement

This research was funded by the National Key R&D Program of China (2024YFC3407800 to H.L. and X.X.), National Natural Scientific Foundation of China (62403390 to P.W.), the Fundamental Research Funds for the Central Universities (XJSJ25016 to P.W.), the Strategic Priority Research Program of the Chinese Academy of Sciences (XDA0460203 to X.X.), the Sponsored by Beijing Nova Program and the National Natural Science Foundation of China (82573343 to X.X.).

## 8 Author contribution

H.D., X.X., P.W., and H.L. supervised the study. H.D., S.M., Y.L., T.T., L.Z., X.X., P.W. and H.L. conceived the project. H.D., S.M. and H.L. designed the method. H.D., S.M., Y.L., J.W., S.Z., P.J. and D.Y. designed and conducted the experiments. All authors prepared the manuscript.

## 9 Ethics declarations

### Competing interests

All authors declare no competing interests.

## Notes

### Competing Interest Statement

The authors have declared no competing interest.

