## Supplementary Files for "SpaTranslator: A deep generative framework for universal spatial multi-omics cross-modality translation"

### A Supplementary Figures

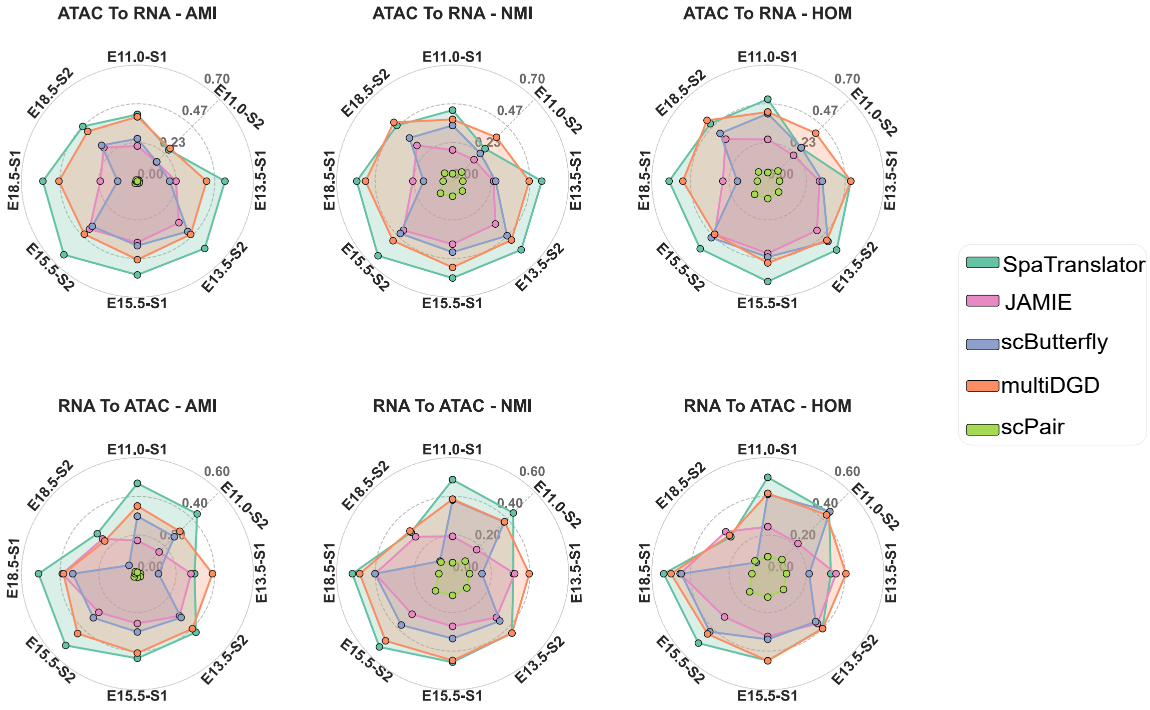


**Fig. S1 The radar charts visualize clustering performance for cross-modality generation tasks on the MISAR-seq dataset.** The upper panel reports three clustering metrics normalized mutual information (NMI), adjusted mutual information (AMI), and homogeneity (HOM) for ATAC-to-RNA translation. Each radar chart compares the performance of five models across eight datasets: E11.0-S1, E11.0-S2, E13.5-S1, E13.5-S2, E15.5-S1, E15.5-S2, E18.5-S1, and E18.5-S2, where E denotes the embryonic timepoint and S denotes the tissue slice. The lower panel presents AMI, NMI, and HOM for RNA-to-ATAC translation.

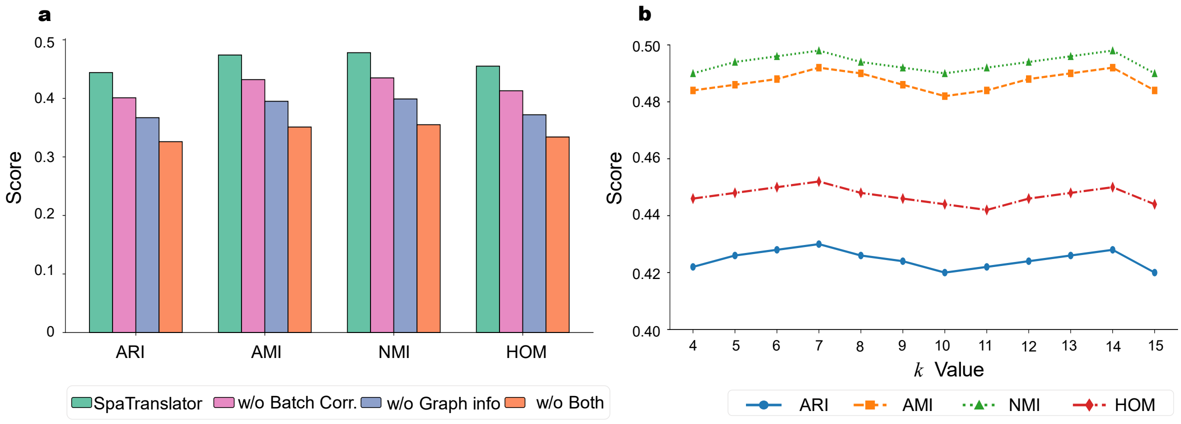


**Fig. S2 Ablation studies of SpaTranslator. a** Clustering performance measured in terms of adjusted Rand index (ARI), normalized mutual information (NMI), adjusted mutual information (AMI), and homogeneity (HOM) with different model components (i.e., batch-correction loss and GNN model) removed. Experiments were conducted on the MISAR-seq dataset. **b** Evaluation of SpaTranslator's robustness to the hyperparameter $k$ in $k$NN graph construction.

### B Supplementary Tables

**Table 1** Summary of spatial multi-omics datasets used in this study.

| Dataset | Tissue/Organ | Technology | # Cells | # Variables | Modality | Slice Name |
|---|---|---|---|---|---|---|
| 1 | Mouse Embryo brain | MISAR-seq | 1243 | 32,285<br>242,024 | RNA<br>ATAC | E11.0-S1 |
| 2 | Mouse Embryo brain | MISAR-seq | 1353 | 32,285<br>213,562 | RNA<br>ATAC | E11.0-S2 |
| 3 | Mouse Embryo brain | MISAR-seq | 1777 | 32,285<br>271,126 | RNA<br>ATAC | E13.5-S1 |
| 4 | Mouse Embryo brain | MISAR-seq | 2183 | 32,285<br>251,555 | RNA<br>ATAC | E13.5-S2 |
| 5 | Mouse Embryo brain | MISAR-seq | 1949 | 32,285<br>265,014 | RNA<br>ATAC | E15.5-S1 |
| 6 | Mouse Embryo brain | MISAR-seq | 1939 | 32,285<br>244,394 | RNA<br>ATAC | E15.5-S2 |
| 7 | Mouse Embryo brain | MISAR-seq | 2091 | 32,285<br>294,734 | RNA<br>ATAC | E18.5-S1 |
| 8 | Mouse Embryo brain | MISAR-seq | 2248 | 32,285<br>223,333 | RNA<br>ATAC | E18.5-S2 |
| 9 | Mouse Embryo brain | spatial ATAC-RNA-seq | 9821 | 23,788<br>121,365 | RNA<br>ATAC | S1 |
| 10 | Mouse Embryo brain | spatial CUT&Tag-RNA-seq | 9752 | 25,881<br>113,273 | RNA<br>H3K27me3 | S2 |
| 11 | Mouse Embryo brain | spatial CUT&Tag-RNA-seq | 9548 | 22,731<br>36,778 | RNA<br>H3K4me3 | S3 |
| 12 | Mouse Embryo brain | spatial CUT&Tag-RNA-seq | 9370 | 23,415<br>125,158 | RNA<br>H3K27ac | S4 |
| 13 | Human Tonsil | 10x Visium coprofiling | 4326 | 18,085<br>31 | RNA<br>ADT | S1 |
| 14 | Human Tonsil | 10x Visium coprofiling | 4519 | 18,085<br>31 | RNA<br>ADT | S2 |
| 15 | Lymph Node | 10x Visium coprofiling | 3484 | 18,085<br>31 | RNA<br>ADT | S1 |
| 16 | Lymph Node | 10x Visium coprofiling | 3359 | 18,085<br>31 | RNA<br>ADT | S2 |

**Table 2** Comparison of different cross-modality generation tools

| Item | SpaTranslator | JAMIE | scButterfly | multiDGD | scPair |
|---|---|---|---|---|---|
| **Data Input** | | | | | |
| Can handle spatial data? | ✅ | ❌ | ❌ | ❌ | ❌ |
| **Strategy** | | | | | |
| Use batch-correction | ✅ | ❌ | ❌ | ❌ | ❌ |
| **Analysis** | | | | | |
| Clustering | ✅ | ✅ | ✅ | ✅ | ✅ |
| Marker gene/protein expression | ✅ | ❌ | ✅ | ❌ | ❌ |
| (For ATAC) Motif analysis | ✅ | ❌ | ❌ | ❌ | ✅ |
| (Multi-omics) GRN inference | ✅ | ❌ | ❌ | ✅ | ❌ |
| **Extendibility** | | | | | |
| RNA-ATAC | ✅ | ✅ | ✅ | ✅ | ✅ |
| RNA-Protein | ✅ | ❌ | ✅ | ❌ | ❌ |

#### C Supplementary Notes: Clustering Metric Definitions

In our quantitative experiments, we employed four widely used clustering evaluation metrics to assess the performance of cross-modality generation. Let $X$ represent the cluster labels predicted by the model and $Y$ represent the ground truth labels. The definitions and formulas for each metric are as follows:

**1. Mutual Information (MI)** MI quantifies the amount of information obtained about one clustering from the other.

$$\mathrm{MI}(X, Y) = \sum_{i=1}^{|X|} \sum_{j=1}^{|Y|} \frac{n_{ij}}{n} \log\left(\frac{n \cdot n_{ij}}{a_i \cdot b_j}\right) \tag{17}$$

where $n_{ij}$ is the number of slices in both predicted cluster $i$ and true cluster $j$, $a_i = \sum_j n_{ij}$, $b_j = \sum_i n_{ij}$, and $n$ is the total number of slices.

**2. Normalized Mutual Information (NMI)** NMI scales MI to the range [0, 1], accounting for cluster size and entropy.

$$\mathrm{NMI}(X, Y) = \frac{2 \cdot \mathrm{MI}(X, Y)}{H(X) + H(Y)} \tag{18}$$

where $H(X) = -\sum_i \frac{a_i}{n} \log\left(\frac{a_i}{n}\right)$ is the entropy of predicted clusters.

**3. Adjusted Mutual Information (AMI)** AMI adjusts MI for chance, accounting for expected mutual information under random assignments.

$$\mathrm{AMI} = \frac{\mathrm{MI}(X, Y) - \mathbb{E}[\mathrm{MI}(X, Y)]}{\max(H(X), H(Y)) - \mathbb{E}[\mathrm{MI}(X, Y)]} \tag{19}$$

**4. Adjusted Rand Index (ARI)** ARI evaluates the similarity of two assignments while correcting for chance grouping.

$$\mathrm{ARI} = \frac{\sum_{ij} \binom{n_{ij}}{2} - \left[\sum_i \binom{a_i}{2} \sum_j \binom{b_j}{2} \Big/ \binom{n}{2}\right]}{\frac{1}{2}\left[\sum_i \binom{a_i}{2} + \sum_j \binom{b_j}{2}\right] - \left[\sum_i \binom{a_i}{2} \sum_j \binom{b_j}{2} \Big/ \binom{n}{2}\right]} \tag{20}$$

**5. Homogeneity (HOM)** Homogeneity assesses if each predicted cluster contains only members of a single true class. It is 1 if all clusters are perfectly pure.

$$\mathrm{HOM}(X, Y) = 1 - \frac{H(Y|X)}{H(Y)} \quad \text{if } H(Y) \neq 0; \text{ otherwise, HOM} = 1 \tag{21}$$

where $H(Y|X)$ is the conditional entropy of the true classes $Y$ given the predicted clusters $X$:

$$H(Y|X) = -\sum_{i=1}^{|X|} \sum_{j=1}^{|Y|} \frac{n_{ij}}{n} \log\left(\frac{n_{ij}}{a_i}\right) \quad \text{if } a_i \neq 0 \tag{22}$$

and $H(Y)$ is the entropy of the true classes $Y$:

$$H(Y) = -\sum_{j=1}^{|Y|} \frac{b_j}{n} \log\left(\frac{b_j}{n}\right) \tag{23}$$

Here, $n_{ij}$ is count of slices in predicted cluster $i$ and true class $j$; $a_i = \sum_j n_{ij}$ (slices in predicted cluster $i$); $b_j = \sum_i n_{ij}$ (slices in true class $j$); $n$ is total slices. Logarithm is natural log. If $a_i = 0$, the term involving $\log(n_{ij}/a_i)$ is 0.

All logarithms are natural logarithms unless otherwise specified. Together, these metrics offer a comprehensive view of clustering performance across multiple statistical perspectives.
